# Shared Structural Features of Miro Binding Control Mitochondrial Homeostasis

**DOI:** 10.1101/2023.07.25.550462

**Authors:** Christian Covill-Cooke, Brian Kwizera, Guillermo López-Doménech, Caleb O.D. Thompson, Ngaam J. Cheung, Ema Cerezo, Martin Peterka, Josef T. Kittler, Benoît Kornmann

## Abstract

Miro proteins are universally conserved mitochondrial calcium-binding GTPases that regulate a multitude of mitochondrial processes, including transport, clearance and lipid trafficking. Miro binds a variety of client proteins involved in these functions. How this binding is operated at the molecular level and whether and how it is important for mitochondrial health, however, remains unknown. Here, we show that known Miro clients all use a similar short motif to bind the same structural element: a highly conserved hydrophobic pocket in the calcium-binding domain of Miro. Using these Miro-binding motifs, we identified direct interactors *de novo*, including yeast Mdm34, and mammalian MTFR1/2/1L, VPS13D and Parkin. Given the shared binding mechanism and conservation across eukaryotes, we propose that Miro is a universal mitochondrial adaptor coordinating mitochondrial health.

**One-Sentence Summary:** Functionally diverse mitochondrial proteins interact with a conserved hydrophobic pocket on the calcium-binding Miro-GTPases.

## Main Text

Mitochondrial function is tightly modulated by homeostatic mechanisms affecting their position, morphology, turnover, and protein and lipid composition. One highly conserved protein family appears central for mitochondrial function, the calcium (Ca2+)-binding Miro-GTPases. Miro binds and often recruits to mitochondria an array of client proteins that are effectors of all of the above processes. These include cytoskeletal adaptors (Trak (*1*–*3*), CENPF (*4*–*6*) and MYO19 (*7*–*9*)), lipid transport contact-site factors (the ER-Mitochondria Encounter Structure, ERMES in yeast (*10, 11*) and VPS13D in metazoans (*12*)) and the mitochondrial quality control E3-ubiquitin ligase Parkin (*13*–*15*) which degrades Miro upon mitochondria-specific autophagy induction (*16*), the failure of which is a hallmark of both idiopathic and familial Parkinson’s disease (PD) (*17*). The functional diversity of Miro clients raises important questions: how does Miro accommodate binding to so many clients? Do they bind simultaneously as (a) large complex(es) or successively through competitive processes? And what significance does this have for the coordination of organelle homeostasis? Miro comprises two GTPase domains (GTPase1 and 2) flanking two Ca2+-binding EF hand with LM helices (ELM1 and 2) (Fig. 1A) (*18*–*20*). Structural information has been gathered on all these domains (*19, 20*), but how Miro binds its partners at the structural level is unexplored.

**Fig. 1.**
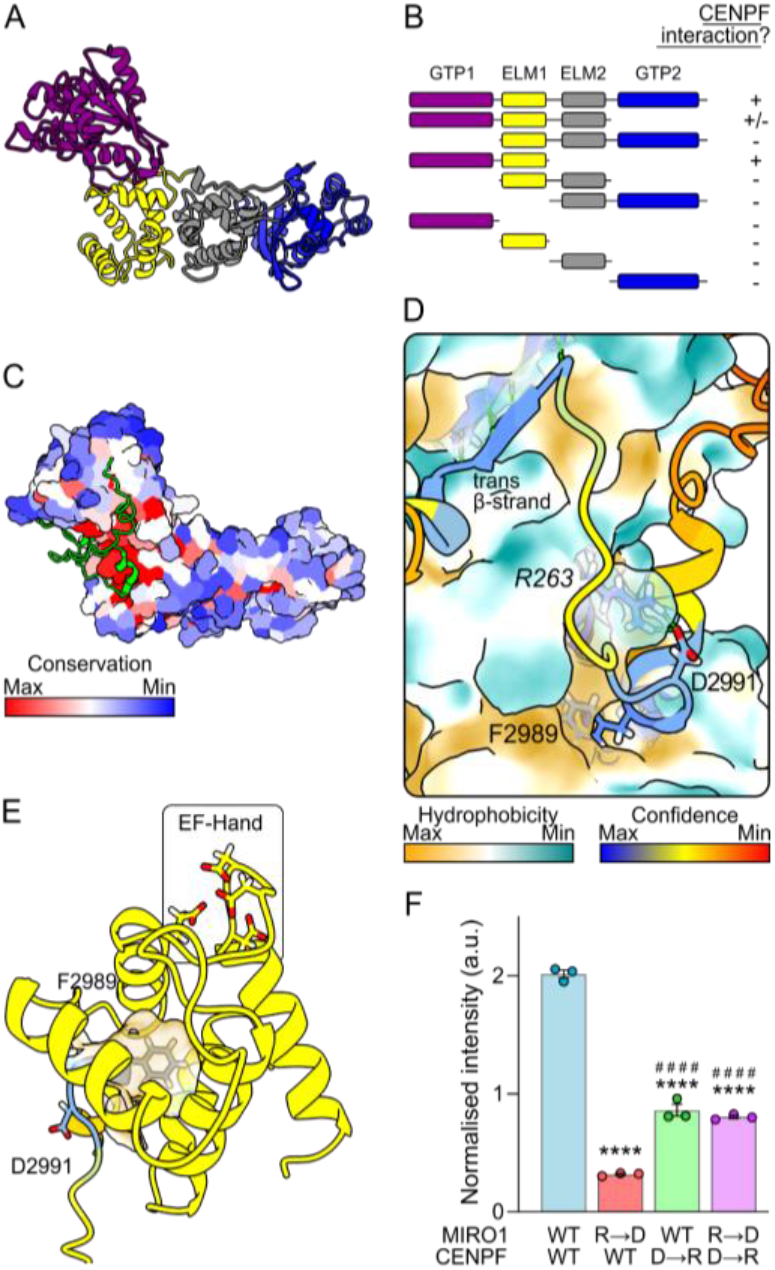
CENPF binds to a conserved hydrophobic pocket in ELM1 of Miro. (**A**) AlphaFold2 predicted structure of human MIRO1 with domains color-coded: purple – GTPase1, yellow – ELM1, grey – ELM2 & blue – GTPase2. The C-terminal transmembrane domain has been removed. (**B**) Schematic showing which truncation constructs of human MIRO1 (prey) bind CENPF-42 (bait) in a yeast two-hybrid assay. + means an interaction was observed; - means no interaction was observed. (**C**) AlphaFold2 multimer prediction of CENPF-42 (shown in green) and human MIRO1. MIRO1 is color-coded according to amino-acid conservation. (**D**) Zoom into the structure in (**C**). Color coding is by prediction confidence for cartoon and by hydrophobicity for MIRO1’s surface. Italicized residues correspond to MIRO1 and non-italicized correspond to CENPF. (**E**) Structural features of the ELF pocket of MIRO1 (yellow) with inserted CENPF-F2989 (color coded as in D) (**F**) Fluorescent yeast two-hybrid assay of wild-type MIRO1 or R263D mutant (R→D), and wild-type CENPF-42 or D2991R mutant (D→R), n=3. Statistical significance was calculated by one-way ANOVA with a Tukey post-hoc test. **** and #### denote p<0.0001 in comparison to WT-MIRO1 + WT-CENPF and WT-MIRO1 + CENPF-D→R, respectively. Graph shows mean ± SEM.

### Identification of a hydrophobic client-binding pocket in Miro

To address these questions, we focused on CENPF on account of its well-defined Miro-binding domain; namely, 42 amino acids within CENPF C-terminus (CENPF-2977-3020, hereafter CENPF-42) necessary and sufficient for direct Miro binding (*4, 5*). To address which Miro domain binds CENPF-42, MIRO1 truncations were generated and cloned into a yeast two-hybrid (Y2H) system with CENPF-42 as bait. We found that both the GTPase1 and ELM1 domains together were necessary and sufficient for CENPF-42 binding (Fig. 1B; Fig. S1). To understand the exact nature of binding, we used the AlphaFold2 multimer model with CENPF-42 and MIRO1 (*21*). AlphaFold2 predicted with high confidence that CENPF binds to MIRO1 at a highly conserved patch (Fig. 1C-D). F2989 - a key phenylalanine residue previously shown to be essential for Miro binding *in vitro* and *in vivo* (*5*) - inserts extensively into a hydrophobic pocket within MIRO1-ELM1 (Fig. 1D-E), opposite to the Ca2+-binding EF-hand, which we call ELM1-domain Leucine-or Phenylalanine-binding (ELF) pocket. Alongside F2989, a conserved aspartate residue (D2991) is predicted to salt-bridge with the conserved R263 on MIRO1 (Fig. 1D). In addition to these ELF-interacting features, a β-strand downstream of F2989 in CENPF (ILR; 3001-3003) makes an antiparallel β-sheet with a β-strand (IETCVE; 141-146) within MIRO1-GTPase1. To validate the AlphaFold2 prediction, we focused on the salt bridge formed by negatively charged CENPF-D2991 and positively charged MIRO1-R263. Using a quantitative fluorescence yeast-two hybrid assay (f-Y2H) as a readout for interaction, we found that mutating MIRO1-R263 to D reduces the interaction. This can be partially rescued by simultaneously mutating CENPF-D2991 to R resulting in a charge swap (Fig. 1F), thus confirming the interaction predicted by AlphaFold2. The CENPF-D2991R mutation alone had comparatively little effect on binding perhaps because MIRO1-R263 can establish compensating bonds with backbone oxygens (see below).

### ELF binding is shared with other Miro interactors

We next sought to understand whether other known interactors bind Miro with a similar conformation. Specific regions of the microtubule motor adaptor proteins, Trak1 and Trak2 (Milton in *Drosophila*), and of the myosin motor MYO19 have been shown to interact with Miro (*1, 2, 7, 8*) (residues 476-700 of mouse Trak2 (*3*) and 898-970 of human MYO19 (*9*)). Therefore, we predicted the interaction of either Trak1, Trak2 or MYO19 Miro-binding domains with MIRO1 in AlphaFold2. All three proteins appear to interact via MIRO1-ELF pocket, with Trak1-L597, Trak2-L581 and MYO19-F948 inserting into the pocket (Fig. 2A-B; Fig. S2A). Both Trak and MYO19 interacting residues show very high conservation (Fig. S2B-C). Indeed, Milton and *Drosophila* Miro (dMiro) are also predicted to interact via the same mechanism (Fig. S2A). 50 amino acid stretches around the pocket-interacting leucine/phenylalanine of mouse-Trak1 and human-MYO19 interacted with MIRO1 in a f-Y2H, with Trak1-L594A (mouse protein) and MYO19-F948A, point mutants abolishing the interaction (Fig. 2C-D), supporting the AlphaFold2 prediction.

**Fig. 2.**
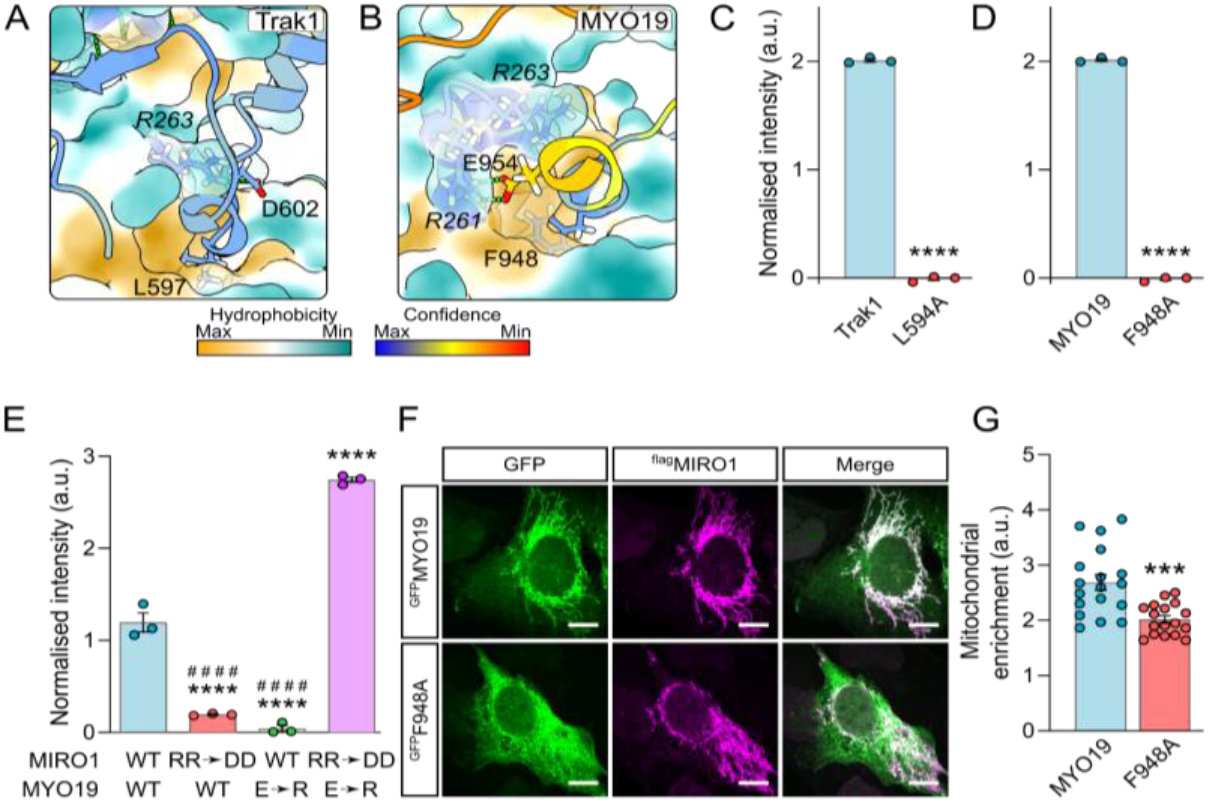
Trak1 and MYO19 bind to the ELF pocket of MIRO1. (**A** and **B**) AlphaFold2 multimer predictions of MIRO1 (surface) with Trak1 and MYO19 (colored as in Fig. 1D), respectively. Italicized residues correspond to MIRO1. (**C** and **D**) Fluorescent yeast two-hybrid assays of human MIRO1 with mouse Trak1-577-620 and human MYO19-919-970, n=3. (**E**) Fluorescent yeast two-hybrid assay of wild-type and R261R-R263D mutant (RR→DD) MIRO1 and wild-type or E954R (E→R) mutant MYO19-917-970, n=3. (**F**) Representative images of U2OS cells transfected with ^flag^MIRO1 (magenta) and either wild-type or F948A ^GFP^MYO19 (green). Scale bars represents 10 μm. (**G**) Quantification of the ratio of mean intensity of ^GFP^MYO19 signal overlapping with ^flag^MIRO1 over non-mitochondrial ^GFP^MYO19 signal. N=18 cells from three independent experiments. (**C, D** and **G**) Statistical significance was calculated by unpaired Student’s t-test. (**E**) Statistical significance was calculated by one-way ANOVA with a Tukey post-hoc test. *** is p<0.001; **** is p<0.0001 in comparison to WT conditions. #### in (**E**) denotes p<0.0001 in comparison to MIRO1-R→D and MYO19-E→R. All graphs show mean ± SEM.

Like for CENPF, MIRO1-R263 was predicted to salt-bridge with either Trak1-D599 or with oxygens in the backbone (Trak2). Accordingly, the MIRO1-R263D mutation reduced binding to Trak1, while Trak1-D599R had little effect (Fig. S2D). In contrast to CENPF though, a charge swap did not rescue interaction, likely because the partial salt bridges made by MIRO1-R263 with Trak1’s backbone are important. We could, however, validate MYO19-binding interface using a charge swap. MYO19-E954 was predicted to make a salt bridge with MIRO1-R261, instead of R263 (Fig. 2B). Yet, while MYO19-E954R mutation substantially reduced interaction with wild-type MIRO1, neither single mutant MIRO1 variants (R261D or R263D) significantly affected binding (Fig. 2E; Fig. S2E). A double MIRO1-R261D-R263D mutant, however, impaired binding. The arginines in the dMiro crystal structure which correspond to human R261 and R263 are not resolved (*19*), suggesting that flexibility in these residues’ orientation accommodates various salt bridges. Charge swapping (i.e., expressing both MYO19-E954R and MIRO1-R261D-R263D) not only rescued, but significantly increased binding, (Fig. 2E), validating the predicted binding conformation.

To assess the relevance of these findings *in vivo*, we took advantage of the fact that the recruitment of MYO19 to mitochondria is partially dependent on Miro (*8*). Overexpression of MIRO1 led to a robust mitochondrial recruitment of MYO19, but not MYO19-F948A, which predominantly localized to the cytoplasm (Fig. 2F-G). A small amount of MYO19-F948A was recruited on mitochondria (Fig. 2F), likely due to the presence of features within the MYO19 C-terminus that allow mitochondrial localization independent of Miro (*9*). Consistent with the fact that Trak1 and Trak2 localize to mitochondria independently of Miro (*8*), Trak1-L594A localized to mitochondria (Fig. S2F). We, therefore, find that the Trak proteins and MYO19 associate with Miro via a shared conserved binding pocket.

### A motif search identifies MTFR1/2/1L as Miro interactors

The identification of a shared mechanism of binding between CENPF, Trak1/2 and MYO19 to Miro raised the possibility that other proteins could interact with the Miro-ELF pocket. To explore this idea, we searched the mitochondrial proteome (MitoCarta3.0) (*22*) for a motif (FADI) based on the ELF binding motif of CENPF. Of five candidates, we focused on MTFR2. MTFR2 is paralogous to MTFR1 and MTFR1L (*23*): two mitochondrial proteins, which also have highly conserved potential Miro binding motifs (MTFR1: FADV; MTFR2: FADI & MTFR1L: LADI) (Fig. S3A). All three proteins were predicted by AlphaFold2 to bind to MIRO1 via the ELF pocket using either a phenylalanine (MTFR1 and MTFR2) or leucine (MTFR1L) residues (Fig. 3A), and in all three cases, these interactions were confirmed using Y2Hs of full-length proteins (Fig. S3B). Mutating the leucine or phenylalanine (Mtfr1-F76A, Mtfr2-F93A, mouse homologues, and MTFR1L-L62A, human homologue) reduced the interaction with MIRO1 (Fig. S3B). Despite all proteins having an acidic residue near the Miro binding motif, none were predicted to make a salt bridge with MIRO1. They were, however, predicted to make a β-sheet (MTFR1: ARL, 91-93; MTFR2: LRF, 91-93, MTFR1L: ARV, 77-79 in human sequences) with MIRO1-GTPase1 (Fig. 3A), like CENPF and Trak.

**Fig. 3.**
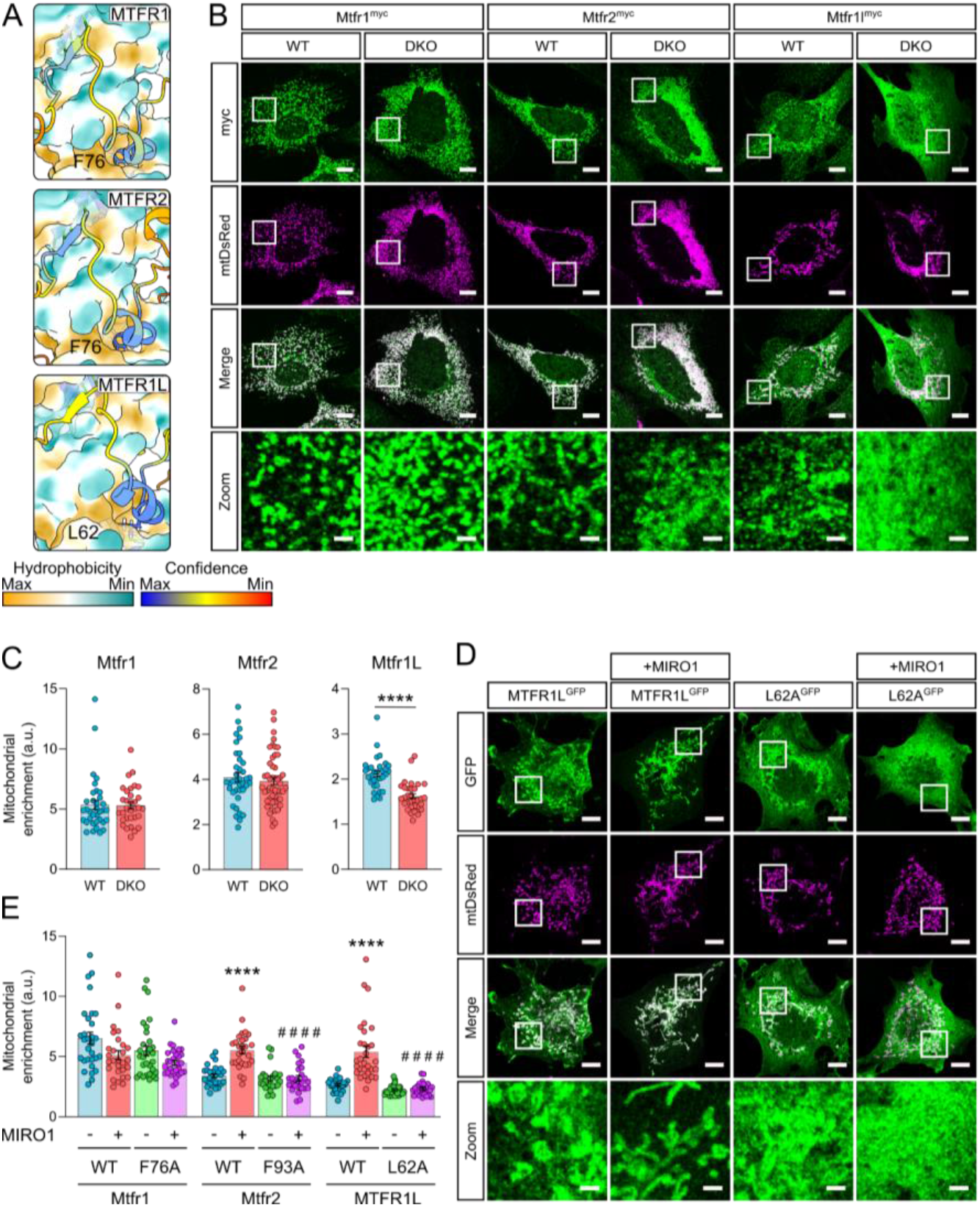
MTFR1/2/1L as novel Miro interactors. (**A**) AlphaFold2 predictions of MTFR1, MTFR2 and MTFR1L and MIRO1 (colored as in Fig. 1D). (**B**) Representative images of myc-tagged mouse Mtfr1, Mtfr2 and Mtfr1l (green) in wild-type and Miro1/2 double knockout mouse embryonic fibroblasts. Mitochondria are stained with mtDsRed (magenta). (**C**) Quantification of mitochondrial localisation of myc-tagged Mtfr1, Mtfr2 and Mtfr1l by calculating the ratio of mean intensity on the mitochondria over non-mitochondrial signal. N=32-49 cells over five independent experiments. Statistical significance was calculated by unpaired Student’s t-test. (**D**) Representative images of wild-type and L62A MTFR1L^GFP^ (green) in Cos7 cells transfected with and without ^myc^MIRO1. Mitochondrial are stained with mtDsRed (magenta). (**E**) Quantification of mitochondrial localisation of wild-type and point mutant Mtfr1, Mtfr2 and MTFR1L, both with and without MIRO1 overexpression, by calculating the ratio of mean intensity on and off the mitochondria. Statistical significance was calculated by one-way ANOVA with post-hoc Tukey test. (**B** and **D**) Scale bars represents 10 μm and 2 μm in zooms. **** is p<0.0001 in comparison to WT conditions. #### is p<0.0001 in comparison to WT+MIRO1. All data are shown as mean ± SEM.

MTFR1, MTFR2 and MTFR1L localize to mitochondria (*23*–*25*). To study if Miro was required for mitochondrial localization, mouse Mtfr1, Mtfr2 and Mtfr1l constructs were expressed in wild-type (WT) and Miro1/2 double knockout mouse embryonic fibroblasts (DKO MEFs). While Mtfr1 and Mtfr2 localized similarly to mitochondria in WT and DKO MEFs, and caused mitochondrial fragmentation as previously described, Mtfr1l mitochondrial localization was reduced upon loss of Miro (Fig. 3B-C). To confirm the role of the Miro-binding motifs *in vivo*, we assessed the recruitment of Mtfr1-F76A, Mtfr2-F93A and MTFR1L-L62A mutants in Cos7 cells. In agreement with the DKO MEF microscopy data, Mtfr1 and Mtfr1-F76A localized to mitochondria, regardless of MIRO1 overexpression (Fig. 3E; Fig. S3C). In contrast, MIRO1 overexpression caused increased recruitment of WT but not of the F93A Mtfr2 mutant (Fig. 3E; Fig. S3D). Similarly, MTFR1L-L62A was not recruited to mitochondria by MIRO1 overexpression, in agreement with the DKO MEFs data (Fig. 3D-E). Therefore, all three MTFR proteins interact with Miro, two of which depend at least partially on Miro for mitochondrial localization. To sum up, a motif search identified novel clients which use the Miro-ELF pocket.

### Conservation of ELF pocket binding

The high conservation of the Miro-ELF pocket (Fig. 1C) and the varied interactors which bind it suggest conservation of this Miro-binding mechanism. We therefore set out to test if non-metazoan Miro orthologues also have this mechanism. Gem1 (Miro orthologue in *Saccharomyces cerevisiae*) is part of ERMES (*10*), a protein complex made up additionally of Mmm1, Mdm12, Mdm34 and Mdm10, that tethers the ER to mitochondria, allows efficient lipid transport between the two compartments, and is essential for tubular mitochondrial morphology (*26, 27*). How Gem1 interacts with other ERMES components is not currently known. By testing each of them, in AlphaFold2, we identified a disordered loop in Mdm34 as interacting with Gem1 (Fig. 4A-B; Fig. S4A). Importantly, this interaction was via a leucine residue (L248) inserting into the cognate ELF pocket of Gem1. An additional salt bridge is present but different from those found in metazoans, and involving Gem1-E242, a residue that is universally conserved, except in metazoans, highlighting divergent evolution. To test if L248 in Mdm34 is required for Gem1 interaction with ERMES, we took advantage of the fact that Gem1 colocalizes in puncta with Mdm34 at ER-mitochondria contacts *(10)*(Fig. 4C). Mutating Mdm34-L248 to alanine in the endogenous locus caused a complete dissociation of Gem1 from ERMES, resulting in a diffuse signal throughout mitochondria (Fig. 4C-D). Importantly, Mdm34-L248A formed foci and mitochondria remained tubular in this condition, indicating that ERMES function wasn’t abolished. Therefore, ERMES binding to Miro’s fungal orthologue is structurally similar to Miro-clients in metazoans. This emphasizes the conservation of the ELF pocket across eukaryotes.

**Figure 4.**
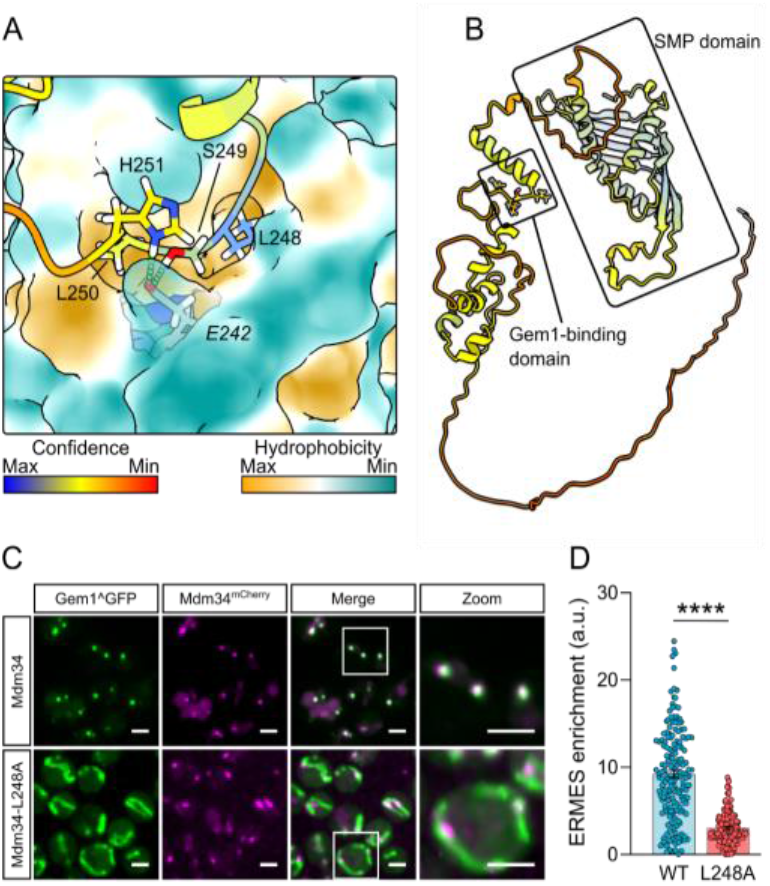
Mdm34-L248 interacts with ELF pocket of Gem1. (**A**) Structural prediction of Saccharomyces cerevisiae Gem1 (surface) with Mdm34 (colored as in Fig. 1D). Italicized residue corresponds to Gem1. (**B**) AlphaFold2 predicted structure of Mdm34 highlighting the lipid-transporting SMP-domain and Gem1-binding domain. (**C**) Representative images of internally GFP-tagged Gem1 (green) in wild-type and Mdm34-L248A budding yeast. Mdm34 was tagged with a C-terminal mCherry (magenta). Scale bars depict 2 μm. (**D**) Quantification of extent of Gem1^GFP colocalizing with Mdm34-mCherry. N=168 cells over two independent experiments. Statistical significance was calculated by unpaired Student’s t-test. **** is p<0.0001. Data are shown as mean ± SEM.

### Parkin and VPS13D bind the Miro-ELF pocket

Having identified a shared binding mechanism for several clients, we could assemble criteria to define binding motifs: i) a conserved phenylalanine or leucine is required for pocket insertion; ii) in mammals, at least, the F/L is often alongside an acidic residue; and iii) the pocket-associating residues are in a conserved disordered loop. Using this knowledge, we set out to identify Miro-binding motifs in other proteins that associate with Miro, focusing on VPS13D and Parkin. VPS13D is a lipid transporter recently described as a Miro interactor which bridges the ER and mitochondria (*12*), is essential in mammals (*28, 29*), and alleles of which cause recessive spinocerebellar ataxia (*30, 31*). VPS13D’s lipid transporting function is homologous to that of ERMES, and yeast Vps13 and ERMES are partially functionally redundant (*32, 33*). Efforts to identify exactly where this interaction occurs on VPS13D and whether it is direct have not been fruitful but a so-called Vps13 adaptor binding (VAB) domain has been proposed (*12*), partly by homology to yeast Vps13, which binds partners through this domain (*33, 34*).

A predicted structure of VPS13D, color-coded by conservation highlighted only two of the many unstructured loops as conserved (Fig. 5A): one comprising the phospho-FFAT motif required for associating to the ER via binding with VAP-A/B (*12*), and the other, we term Miro-Binding Motif (MBM), adjacent to the VAB domain. This second loop contains a conserved L2554 which AlphaFold2 predicted to insert into the ELF pocket (Fig. 5B; Fig. S5). To confirm this prediction, we used a mitochondrial recruitment assay. Overexpression of Miro caused significant recruitment of wild-type VPS13D to mitochondria (*12*) (Fig. 5C-D). Remarkably, the recruitment of VPS13D-L2554A was severely blunted, despite differing from wild-type by a mere 42 Da over ∼492,000 Da. We conclude that VPS13D binding to Miro-ELF pocket is a key part of its association with mitochondria.

**Fig. 5.**
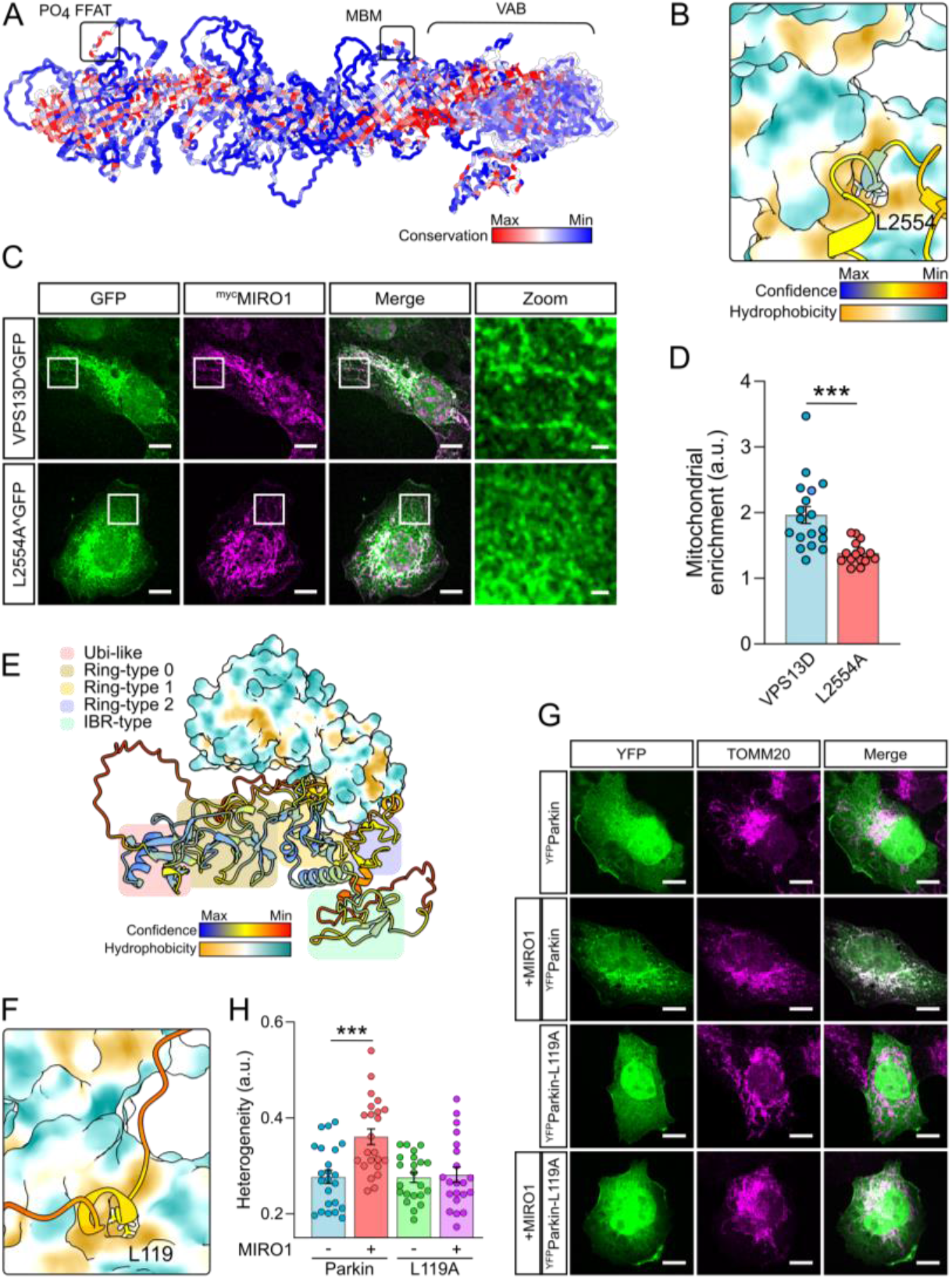
Conserved leucine residues in both VPS13D and Parkin interact with the Miro ELF pocket. (**A**) Predicted full-length structure of human VPS13D with residues colored by conservation. PO4 FFAT = phospho-FFAT motif for VAP binding; MBM = Miro binding motif; VAB = VPS13 adaptor binding domain. (**B**) AlphaFold2 multimer prediction of MIRO1 (surface) and VPS13D. (**C**) Representative images of internally GFP-tagged wild-type and L2554A mutant VPS13D (green) in Cos7 cells overexpressing ^myc^MIRO1 (magenta). (**D**) Quantification of mean mitochondrial intensity divided by mean intensity in cytoplasm. N=15-18 cells over three independent experiments. Statistical significance was calculated by an unpaired Student’s t-test. (**E**) Structural prediction of interaction between MIRO1 (surface) with full-length Parkin. Colored boxes highlight the individual predicted domains of Parkin. (**F**) Zoom of structural predictions of Miro-ELF pocket and Miro-binding motif of Parkin. (**G**) Representative images of wild-type and L119A ^YFP^Parkin (green) in U2OS cells either with or without ^myc^MIRO1 overexpression. Mitochondria were stained with TOMM20 antibody (magenta). (**H**) Quantification of the heterogeneity of YFP signal from wild-type and L119A Parkin, both with and without MIRO1 overexpression. Statistical significance was calculated by one-way ANOVA with Tukey post-hoc test. All data are shown as mean ± SEM. *** is p<0.001. Scale bars represents 10 μm and 2 μm in zooms.

Parkin rapidly ubiquitinates Miro during mitochondrial damage, as part of mitophagy (*16*). Miro overexpression increases Parkin recruitment to mitochondria irrespective of mitochondrial damage (*13*–*15*), but whether this is due to direct interaction is not known. An AlphaFold2 prediction of full-length Parkin with MIRO1 suggested that Parkin might bind to the ELF pocket using the conserved L119 (Fig. 5E-F; Fig. S5). To validate this prediction, we imaged WT and L119A mutant Parkin. We observed partial wild-type Parkin recruitment onto mitochondria upon MIRO1 overexpression (Fig. 5G-H), which we quantified as previously (*14*), as an increase in signal heterogeneity. Importantly, Parkin-L119A staining remained homogenously cytosolic even upon MIRO1 overexpression, highlighting this leucine being critical for Miro-Parkin interaction and supporting Parkin binding Miro-ELF pocket directly (Fig. 5G-H).

## Discussion

### The various biochemical features affecting Miro binding

Here, we identify that Miro proteins interact with a variety of partners with a similar conformation, whereby interactors bind a hydrophobic pocket. All predictions performed with MIRO1 were performed with MIRO2, and no differences were observed. Previous work has suggested client binding is dependent on Miro’s calcium and nucleotide status. The ELF pocket is made in part by the Ca2+-binding EF-hand. Interestingly, Miro’s crystal structure shows a density corresponding to an unknown small-molecule ligand occupying the ELF pocket, enlarging it, thus preventing Ca2+ binding (*19*). Whether this corresponds to a physiological ligand competing partners out of Miro-ELF pocket or is an artifact of protein expression is yet unknown, but these findings suggest that Ca2+, ligand and client binding in the ELF pocket are mutually exclusive.

In addition to the ELF pocket, several clients (CENPF, Trak1/2, MTFR1/2/1L & Mdm34) establish an antiparallel β-strand with Miro-GTPase1. The significance of this feature is difficult to assess as β-sheets do not obviously involve mutable side chains. Nonetheless, the β-strand is established at a highly conserved patch of GTPase1 previously named the SELFYY surface (named after a conserved peptide) (*20*). How nucleotide binding in the GTPase1 domain affects client binding is unclear. For example, clients like VPS13D do not make any contact with the GTPase1 domain, yet their interaction is dependent on a wild-type GTPase1 domain (*12*). It is therefore possible that nucleotide binding elicits larger allosteric changes; for instance, controlling the flexible hinge positioning the GTPase1 domain, which could control access to the ELF pocket.

Although the general Miro-binding conformation is shared, details are intriguingly different (e.g., leucine vs phenylalanine, with or without salt bridge, or β-sheet). For instance, a glycine residue preceding the leucine/phenylalanine is found in several clients. In CENPF, this glycine is vital for binding (*5*). Why it is not important in all clients might boil down to the slightly different conformations taken by the client’s backbone when entering and exiting the ELF pocket. Chains with glycines harbor bond angles that other amino acids cannot adopt (Fig. S6). One explanation for flexibility in the motif is that it is an easily “evolvable” element that can likely be exploited when an interaction with Miro becomes a competitive advantage. A parallel might be drawn to the VAP proteins that bind short and diverse FFAT motifs to recruit proteins and whole organelles to the ER (*35*). As such Miro might be regarded as a general and regulatable adaptor to recruit proteins and organelles to mitochondria.

### Miro proteins as coordinators of mitochondrial homeostasis

The identification of key leucines/phenylalanines in clients provides an opportunity to decipher the importance of their binding to Miro. Single leucine/phenylalanine point mutants provide a means to perturb the residue specifically required for Miro binding whilst keeping the rest intact, i.e., maintaining Miro-independent processes which would be lost with gene deletions. Indeed, we have shown that mutating CENPF-F2989 prevented recruitment to mitochondria, but yielded surprisingly healthy mice (*5*). It will be crucial to assess the phenotypic consequences of specifically disrupting interaction with Miro for other partners as well. For instance, Trak proteins are recruited to mitochondria independently of Miro (*8*). What, therefore, is the specific role of Miro binding in their microtubule-dependent mitochondrial transport function?

A noteworthy consequence of a shared binding site on Miro for its clients, is that the roles of Miro in microtubule-dependent trafficking, actin dynamics, mitochondrial morphology, lipid transport and mitophagy must be competitive, further suggesting that there is, to some extent, competition between these processes themselves. Previous groups have proposed elements of this idea, as in the model where mitochondria must be released from microtubules to be efficiently degraded (*16, 17, 36*) or attached to actin filaments during early-, and microtubule tips during late-mitosis (*37, 38*). This competition might now be traced at the molecular level to competitive binding. This mechanism is likely shared in many eukaryotic species, for a currently unknown number of processes at mitochondria. For instance, we do not know any client for plant Miro. We therefore expect that Miro function is to be a central point at the outer mitochondrial membrane to coordinate mitochondrial homeostasis.

## Acknowledgments

The authors are grateful for manuscript feedback from Agnès Michel and Viktoriya S. Toncheva.

The authors also gratefully acknowledge the Micron Advanced Bioimaging Facility (supported by Wellcome Strategic Awards 091911/B/10/Z and 107457/Z/15/Z) for their support & assistance in this work.

## Funding

Wellcome Trust grant 214291/Z/18/Z (BK)

Wellcome Trust grants 223202/Z/21/Z and 222/519/Z/21/Z (JTK).

MRC PhD studentship 2397875 (CODT)

## Author contributions

Conceptualization: CCC, BK, JTK, GLD, BKw, CODT, EC, NJC, MP

Methodology: CCC, BK, JTK, GLD, BKw, CODT, EC, NJC, MP

Investigation: CCC, BK, GLD, BKw, CODT, EC, NJC, MP

Visualization: CCC, BK

Funding acquisition: BK, JTK

Project administration: BK, JTK

Supervision: CCC, BK, JTK, NJC

Writing – original draft: CCC, BK

Writing – review & editing: CCC, BK, JTK, GLD, BKw, CODT, MP

## Data and materials availability

All data are available in the main text or the supplementary materials.

## Supplementary Materials

### Materials and Methods

#### DNA constructs

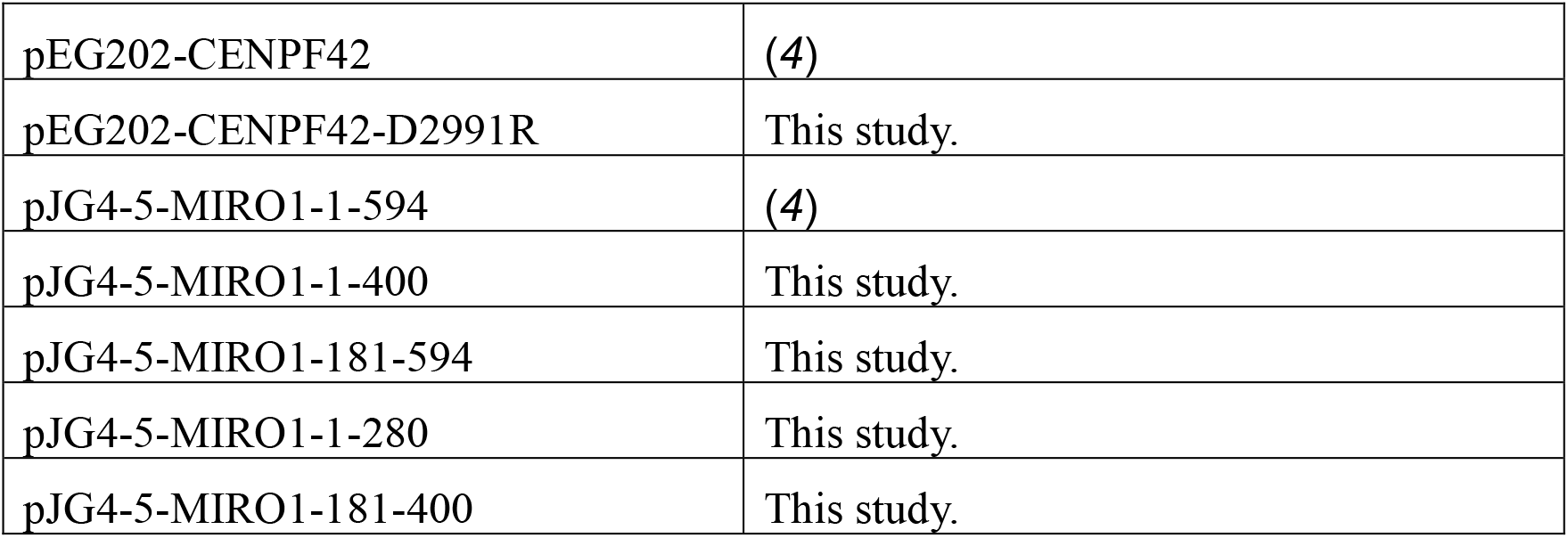
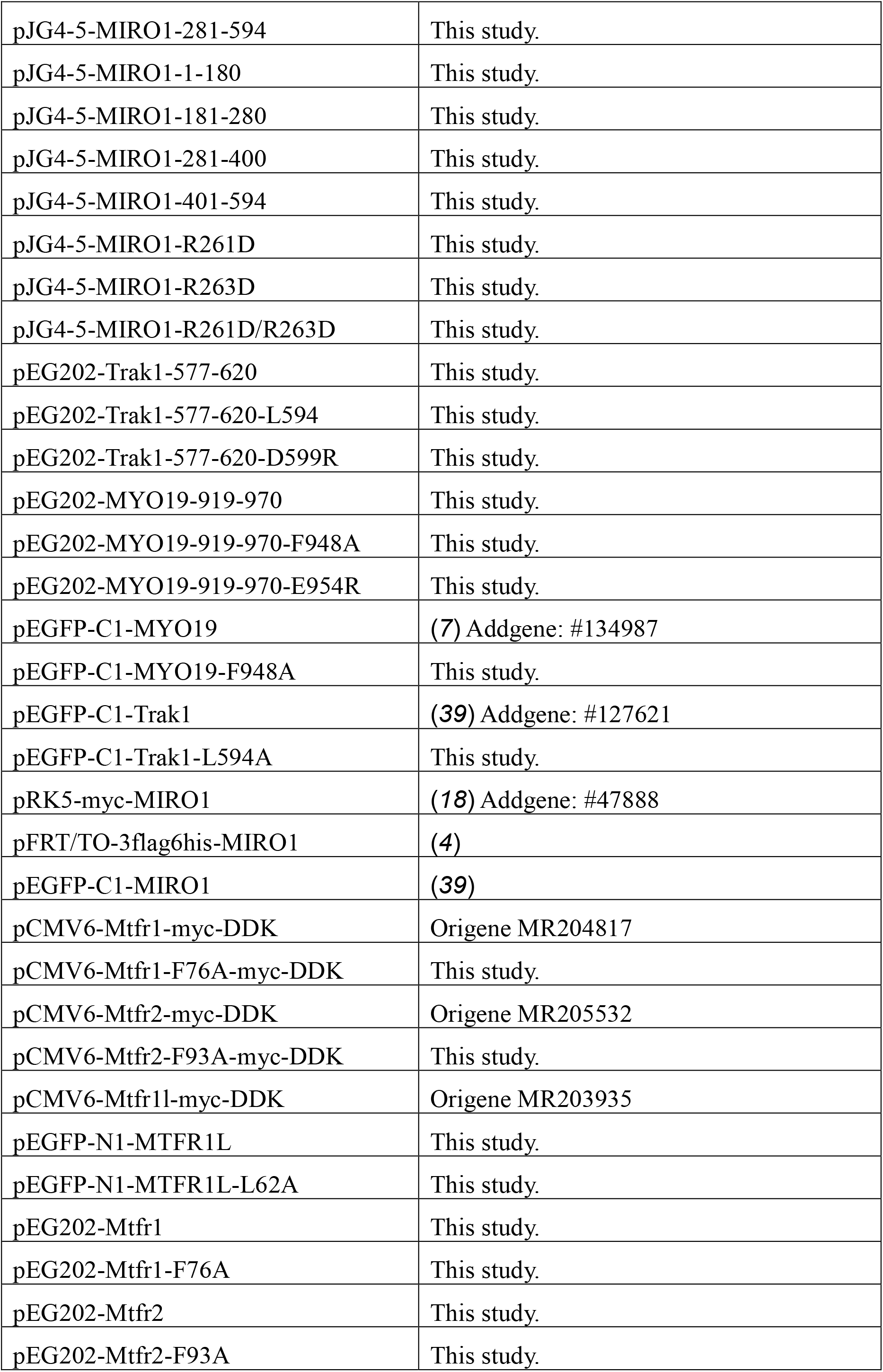
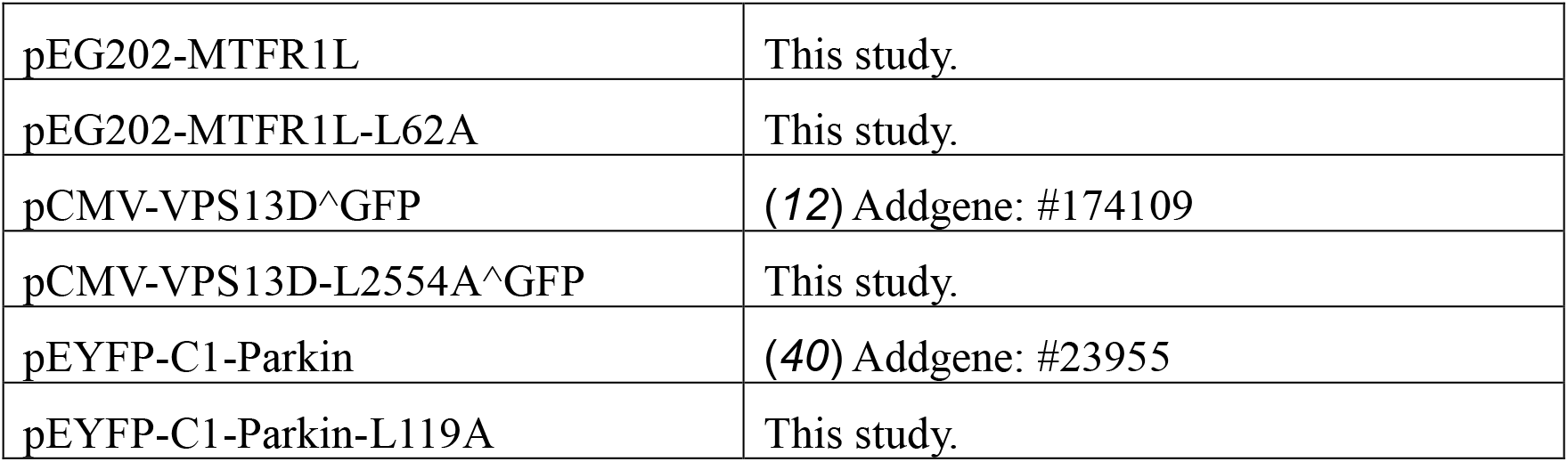

#### Antibodies and dyes

Primary antibodies: mouse anti-myc (9E10 at 1:1,000), mouse anti-Flag (M2 at 1:1,000), rabbit-TOMM20 (Santa Cruz -sc-11415, 1:500), rabbit anti-myc tag antibody (Abcam - ab9106, 1:1,000). Secondary antibodies: donkey anti-mouse IgG H&L-AlexaFluor-647 (Abcam ab1501017 at 1:500), donkey anti-rabbit IgG H&L-AlexaFluor-568 (Abcam ab175470 at 1:500). MitoTracker Orange CMTMRos was obtained from Thermo Fisher Scientific (M7510).

#### Yeast and mammalian cell lines

|  |  |
| --- | --- |
| EGY48 | Yeast-two hybrid competent yeast. |
| ByK302 | By4741 *MDM34*-mCherry::His3; Gem1<sub>263</sub>GFP |
| ByK2029 | ByK302 with L248A mutation in *MDM34* by CRISPR/Cas9 mutagenesis |

##### Generating yeast strains

*MDM34*-mCherry::HIS3 and internally GFP-tagged Gem1 (at position 263) were generated previously (*26, 41*). A L248A mutation in *MDM34* was generated using the CRISPR/Cas9 system (*42*) using the following gRNA: 5’-tttcaagcattgtgtcgtcgagg-3’ and a repair template including the desired mutation.

##### Mammalian cells

U2OS and Cos7 cells were cultured in DMEM with 4.5 g/L glucose plus 10 % fetal bovine serum, GlutaMAX and penicillin/streptomycin. Wild-type and Miro1/2 double knockout mouse-embryonic fibroblasts (MEFs), characterized previously (*8*), were cultured in DMEM with 4.5 g/L glucose plus 15 % fetal bovine serum, GlutaMAX and penicillin/streptomycin. For fixed imaging, MEFs were seeded on fibronectin-coated coverslips.

#### Yeast-two hybrid

All yeast-two hybrid assays were based on *LexA* fusion proteins. EGY48 yeast were transformed with a pJG4-5 MIRO1 construct (prey) and a bait containing plasmid (pEG202). For growth assays, yeasts were streaked on SC-Leu+Gal media and grown at 30°C. For fluorescence yeast-two hybrid assays a modified protocol from (*43*) was used. Yeasts were grown overnight in SC-Trp-His-Ura + 2% raffinose and 0.2% glucose and then switch to overnight in SC-Trp-His-Ura + 2% galactose. The following day, 2,000,000 cells for each condition were collected and resuspended in ice-cold 70% ethanol and shaken at 2,850 rpm for 5 minutes to permeabilized the cells. Cells were then pelleted and resuspended in 10 ml buffer Z (0.06 M Na_2_HPO_4_, 0.04 M NaH_2_PO_4_, 0.01 M KCl, 0.001 M MgSO_4_ and 0.27% 2-mercaptoethanol) for CENPF and Trak1 and 1 ml of buffer Z for MYO19 due to differences in signal intensity. 50 μl of cell suspension and 50 μl of Fluorescein di-beta-D-galactopyranoside (FDG; 0.5 mg/ml dissolved in 98% water, 1% ethanol and 1% DMSO; Stratatech - 14001) were then mixed together and imaged using the Fluorescein-FITC channel on an iBright-FL1500. Data are well fluorescence minus signal for empty vector divided by mean signal over all wells

#### Structural predictions

All structure figures were generated in ChimeraX (*44*).

##### AlphaFold predictions

Monomeric Mdm34 and MIRO1 AlphaFold2 predictions were obtained from the AlphaFold-European Bioinformatics Institute database. Protein-protein interaction predictions were made using the AlphaFold2 multimer model (*21*) - ran both remotely and on the open source AlphaFold.ipynb on Google Colab.

##### VPS13D

A full-length VPS13D structure was predicted using a coarse-grained molecular dynamics simulation (MoDyFing) to fold the 3D structure of VPS13D from its primary sequence depending on the structural constraints (residue-distances and torsion angles) that are inferred by deep learning methods. The torsion angles (phi and psi) were predicted by the ESIDEN tool (*45*), while the distance between pairwise residues was inferred by the ProSpr mode (*46*). The constraints were used to predict protein 3D structure of no more than 900 residues. As such, VPS13D was split into seven fragments including three overlapping fragments. We leveraged the MoDyFing tool to fold each fragment using the inferred constraints and implemented the MODELLER tool to assemble the predicted structures of the seven fragments.

##### Mapping conservation of amino acids

Residue conservation for MIRO1 and VPS13D was made using the top 1,000 conserved sequence in comparison to the human protein.

#### Bioinformatics

Multiple sequence alignments were generated with MUSCLE and displayed with jalview. Ramachandran plots used density data from (*47*).

#### Fluorescence microscopy

##### Live imaging of yeast

Yeast saturated cultures were reseeded to OD of 0.1 in YPD and left to recover for six hours. Roughly 500,000 cells were then washed in SC media and plated on a microscope slide with a coverslip on top. Images were obtained using a IX81 Olympus inverted spinning disk microscope with an EM-CCD camera (Hamamatsu Photonics) using a 100x oil objective (NA=1.4).

##### Fixed imaging of mammalian cells

Cells were fixed with 4% paraformaldehyde in PBS for 10 minutes at room temperature and blocked with 5 mg/ml bovine serum albumin, 10% horse serum and 0.2% Triton X100 diluted in PBS. Primary and secondary antibodies were diluted in blocking buffer and used to stain cells for one hour at room temperature. Images were taken on either a IX81 Olympus inverted spinning disk microscope with an EM-CCD camera (Hamamatsu Photonics) using a 100x oil objective (NA=1.4) or a Zeiss LSM700 confocal using a 63 × oil objective (NA = 1.4).

#### Image analysis

##### Mitochondrial enrichment

Mitochondrial enrichment of fluorescent signal was calculated by dividing the mean fluorescence overlapping with a thresholded mitochondrial marker (e.g., Tom20 or mtDsRed) divided by the mean fluorescence intensity in the rest of the cell. For VPS13D, due to the high intensity of VPS13D^GFP signal at the Golgi, blind analysis was performed on 8 μm^2^ crops of cells at a point where mitochondria are tubular and away from the perinuclear GFP signal.

##### ERMES enrichment

Gem1 enrichment at ERMES was calculated as the integrated density of signal overlapping with Mdm34-mCherry divided by the integrated density of GFP signal in the whole cell. The cell was identified using the YeastMate plugin (*48*) in ImageJ.

##### Heterogeneity of Parkin signal

Data were blinded and quantified by taking 8 μm^2^ crops of cells at a point where mitochondria are tubular and away from the nucleus, using the TOMM20 stain for reference. The coefficient of variation of YFP-Parkin signal was then calculated by dividing the standard deviation of YFP signal intensity by the mean intensity.

## References

1. S. Fransson, A. Ruusala, P. Aspenström, The atypical Rho GTPases Miro-1 and Miro-2 have essential roles in mitochondrial trafficking. Biochem. Biophys. Res. Commun. 344, 500–510 (2006).

2. 2. E. E. Glater, L. J. Megeath, R. S. Stowers, T. L. Schwarz, Axonal transport of mitochondria requires milton to recruit kinesin heavy chain and is light chain independent. J. Cell Biol. 173, 545–557 (2006).

3. A. F. MacAskill, K. Brickley, F. A. Stephenson, J. T. Kittler, GTPase dependent recruitment of Grif-1 by Miro1 regulates mitochondrial trafficking in hippocampal neurons. Mol. Cell. Neurosci. 40, 301–312 (2009).

4. G. Kanfer et al., Mitotic redistribution of the mitochondrial network by Miro and Cenp-F. Nat. Commun. 6, 8015 (2015).

5. M. Peterka, B. Kornmann, Miro-dependent mitochondrial pool of CENP-F and its farnesylated C-terminal domain are dispensable for normal development in mice. PloS Genet. 15, e1008050 (2019).

6. G. Kanfer et al., CENP-F couples cargo to growing and shortening microtubule ends. Mol. Biol. Cell. 28, 2400–2409 (2017).

7. S. J. Oeding et al., Identification of Miro1 and Miro2 as mitochondrial receptors for myosin XIX. J. Cell Sci. 131 (2018), doi:10.1242/jcs.219469.

8. G. López-Doménech et al., Miro proteins coordinate microtubule- and actin-dependent mitochondrial transport and distribution. EMBO J. 37, 321–336 (2018).

9. J. L. Bocanegra et al., The MyMOMA domain of MYO19 encodes for distinct Miro-dependent and Miro-independent mechanisms of interaction with mitochondrial membranes. Cytoskeleton (Hoboken). 77, 149–166 (2020).

10. B. Kornmann, C. Osman, P. Walter, The conserved GTPase Gem1 regulates endoplasmic reticulum-mitochondria connections. Proc. Natl. Acad. Sci. USA. 108, 14151–14156 (2011).

11. D. A. Stroud et al., Composition and topology of the endoplasmic reticulum-mitochondria encounter structure. J. Mol. Biol. 413, 743–750 (2011).

12. 12. A. Guillén-Samander et al., VPS13D bridges the ER to mitochondria and peroxisomes via Miro. J. Cell Biol. 220 (2021), doi:10.1083/jcb.202010004.

13. G. López-Doménech et al., Loss of neuronal Miro1 disrupts mitophagy and induces hyperactivation of the integrated stress response. EMBO J. 40, e100715 (2021).

14. D. Safiulina et al., Miro proteins prime mitochondria for Parkin translocation and mitophagy. EMBO J. 38 (2019), doi:10.15252/embj.201899384.

15. E. Shlevkov, T. Kramer, J. Schapansky, M. J. LaVoie, T. L. Schwarz, Miro phosphorylation sites regulate Parkin recruitment and mitochondrial motility. Proc. Natl. Acad. Sci. USA. 113, E6097–E6106 (2016).

16. X. Wang et al., PINK1 and Parkin target Miro for phosphorylation and degradation to arrest mitochondrial motility. Cell. 147, 893–906 (2011).

17. C.-H. Hsieh et al., Functional impairment in miro degradation and mitophagy is a shared feature in familial and sporadic parkinson’s disease. Cell Stem Cell. 19, 709–724 (2016).

18. A. Fransson, A. Ruusala, P. Aspenström, Atypical Rho GTPases have roles in mitochondrial homeostasis and apoptosis. J. Biol. Chem. 278, 6495–6502 (2003).

19. J. L. Klosowiak et al., Structural coupling of the EF hand and C-terminal GTPase domains in the mitochondrial protein Miro. EMBO Rep. 14, 968–974 (2013).

20. K. P. Smith et al., Insight into human Miro1/2 domain organization based on the structure of its N-terminal GTPase. J. Struct. Biol. 212, 107656 (2020).

21. R. Evans et al., Protein complex prediction with AlphaFold-Multimer. BioRxiv (2021), 10 doi:10.1101/2021.10.04.463034.

22. S. Rath et al., MitoCarta3.0: an updated mitochondrial proteome now with sub-organelle localization and pathway annotations. Nucleic Acids Res. 49, D1541–D1547 (2021).

23. M. Monticone et al., The nuclear genes Mtfr1 and Dufd1 regulate mitochondrial dynamic and cellular respiration. J. Cell Physiol. 225, 767–776 (2010).

24. L. Tilokani et al., AMPK-dependent phosphorylation of MTFR1L regulates mitochondrial morphology. Sci. Adv. 8, eabo7956 (2022).

25. L. Tonachini et al., Chondrocyte protein with a poly-proline region (CHPPR) is a novel mitochondrial protein and promotes mitochondrial fission. J. Cell Physiol. 201, 470–482 (2004).

26. B. Kornmann et al., An ER-mitochondria tethering complex revealed by a synthetic biology screen. Science. 325, 477–481 (2009).

27. A. T. John Peter, C. Petrungaro, M. Peter, B. Kornmann, METALIC reveals interorganelle lipid flux in live cells by enzymatic mass tagging. Nat. Cell Biol. 24, 996–1004 (2022).

28. V. A. Blomen et al., Gene essentiality and synthetic lethality in haploid human cells. Science. 350, 1092–1096 (2015).

29. T. Wang et al., Identification and characterization of essential genes in the human genome. Science. 350, 1096–1101 (2015).

30. E. Seong et al., Mutations in VPS13D lead to a new recessive ataxia with spasticity and mitochondrial defects. Ann. Neurol. 83, 1075–1088 (2018).

31. J. Gauthier et al., Recessive mutations in VPS13D cause childhood onset movement disorders. Ann. Neurol. 83, 1089–1095 (2018).

32. A. B. Lang, A. T. John Peter, P. Walter, B. Kornmann, ER-mitochondrial junctions can be bypassed by dominant mutations in the endosomal protein Vps13. J. Cell Biol. 210, 883– 890 (2015).

33. A. T. John Peter et al., Vps13-Mcp1 interact at vacuole-mitochondria interfaces and bypass ER-mitochondria contact sites. J. Cell Biol. 216, 3219–3229 (2017).

34. B. D. M. Bean et al., Competitive organelle-specific adaptors recruit Vps13 to membrane contact sites. J. Cell Biol. 217, 3593–3607 (2018).

35. S. E. Murphy, T. P. Levine, VAP, a Versatile Access Point for the Endoplasmic Reticulum: Review and analysis of FFAT-like motifs in the VAPome. Biochim. Biophys. Acta. 1861, 952–961 (2016).

36. G. Ashrafi, J. S. Schlehe, M. J. LaVoie, T. L. Schwarz, Mitophagy of damaged mitochondria occurs locally in distal neuronal axons and requires PINK1 and Parkin. J. Cell Biol. 206, 655–670 (2014).

37. J. Y.-M. Chung, J. A. Steen, T. L. Schwarz, Phosphorylation-Induced Motor Shedding Is Required at Mitosis for Proper Distribution and Passive Inheritance of Mitochondria. Cell Rep. 16, 2142–2155 (2016).

38. G. Kanfer, B. Kornmann, Dynamics of the mitochondrial network during mitosis. Biochem. Soc. Trans. 44, 510–516 (2016).

39. N. Birsa et al., Lysine 27 ubiquitination of the mitochondrial transport protein Miro is dependent on serine 65 of the Parkin ubiquitin ligase. J. Biol. Chem. 289, 14569–14582 (2014)

40. D. Narendra, A. Tanaka, D.-F. Suen, R. J. Youle, Parkin is recruited selectively to impaired mitochondria and promotes their autophagy. J. Cell Biol. 183, 795–803 (2008).

41. A. M. English et al., ER-mitochondria contacts promote mitochondrial-derived compartment biogenesis. J. Cell Biol. 219 (2020), doi:10.1083/jcb.202002144.

42. G. Hu, S. Luo, H. Rao, H. Cheng, X. Gan, A Simple PCR-based Strategy for the Introduction of Point Mutations in the Yeast Saccharomyces cerevisiae via CRISPR/Cas9. Biochem. Mol. Biol. J. 4 (2018), doi:10.21767/2471-8084.100058.

43. A. Plovins, A. M. Alvarez, M. Ibañez, M. Molina, C. Nombela, Use of fluorescein-di-beta-D-galactopyranoside (FDG) and C12-FDG as substrates for beta-galactosidase detection by flow cytometry in animal, bacterial, and yeast cells. Appl. Environ. Microbiol. 60, 4638– 25 4641 (1994).

44. E. F. Pettersen et al., UCSF ChimeraX: structure visualization for researchers, educators, and developers. Protein Sci. 30, 70–82 (2021).

45. Y.-C. Xu, T.-J. ShangGuan, X.-M. Ding, N. J. Cheung, Accurate prediction of protein torsion angles using evolutionary signatures and recurrent neural network. Sci. Rep. 11, 30 21033 (2021).

46. J. Stern, B. Hedelius, O. Fisher, W. M. Billings, D. Della Corte, Evaluation of deep neural network prospr for accurate protein distance predictions on CASP14 targets. Int. J. Mol. Sci. 22 (2021), doi:10.3390/ijms222312835.

47. S. C. Lovell et al., Structure validation by Calpha geometry: phi,psi and Cbeta deviation. Proteins. 50, 437–450 (2003).

48. D. Bunk et al., YeastMate: neural network-assisted segmentation of mating and budding events in Saccharomyces cerevisiae. Bioinformatics. 38, 2667–2669 (2022).

