## Supplementary Figures for "Shared Structural Features of Miro Binding Control Mitochondrial Homeostasis"

Supp. Fig. 1

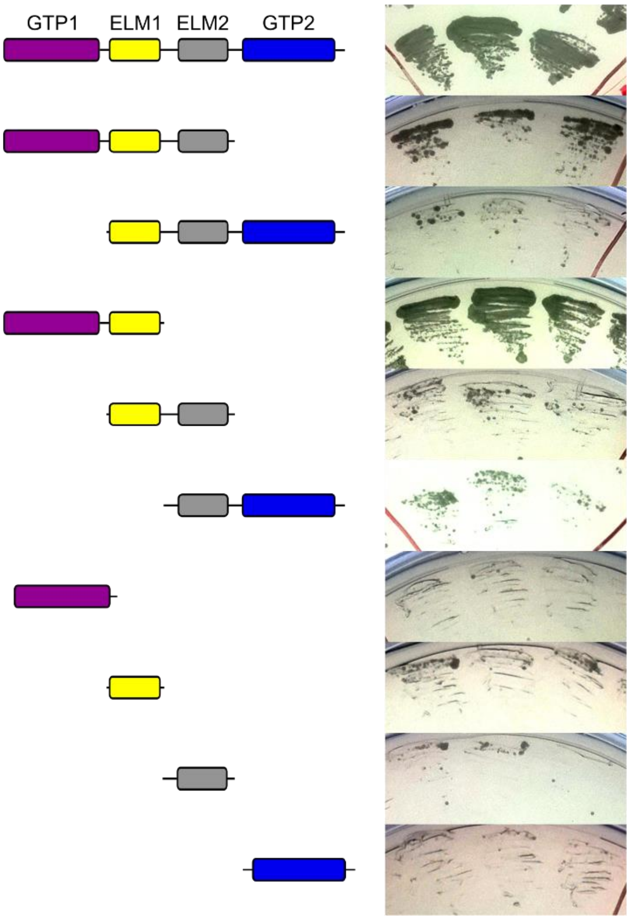


**Fig. S1. GTPase1 and ELM1 of MIRO1 are necessary and sufficient for CENPF binding.** Representative yeast two-hybrid of CENPF-42 (bait) with MIRO1 truncations (prey). Each streak is from an independently generated strain.

Supp. Fig. 2

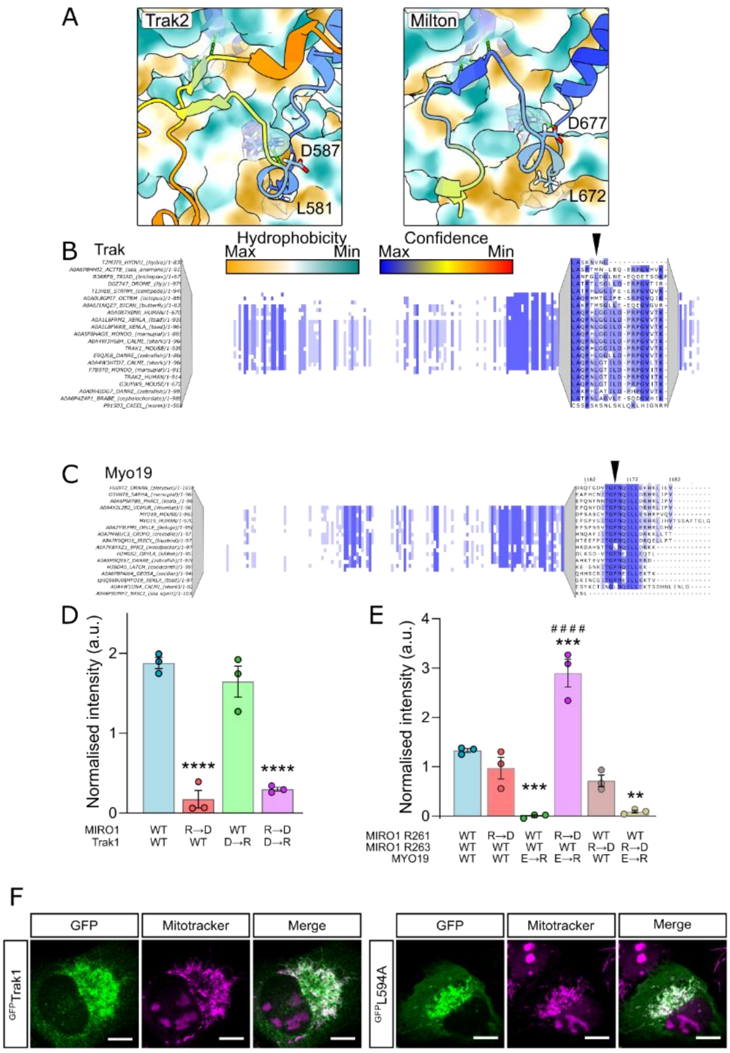


**Fig. S2. Conserved features in Trak1 and MYO19 that interact with MIRO1.** (**A**) AlphaFold2 multimer predictions of human Trak2 with MIRO1 (surface) or *Drosophila melanogaster* Miro (surface) and Milton (Trak orthologue). Color coding as in Fig 1D. (**B**) Sequence alignment of Trak orthologues around the Miro-binding motif. (**C**) Sequence alignment of MYO19 orthologues around the Miro-binding motif. (**D**) Quantification of fluorescence yeast two-hybrid of wild-type and charge swapped mutants of MIRO1 (prey) and mouse Trak1-577-620 (bait). R→D and D→R are MIRO1-R263D and Trak1-D599R, respectively, n=3. (**E**) Quantification of fluorescence yeast two-hybrid of wild-type and charge swapped mutants of MIRO1 (prey) and MYO19-919-970 (bait). R→D and E→R are MIRO1-R261D / MIRO1-R263D and MYO19-E954R, respectively, n=3. (**F**) Representative images of wild-type and L594A mouse $^{GFP}$Trak1 (green) in U2OS cells. Mitochondria are stained with Mitotracker-Orange (magenta). Scale bars represents 10 µm. (**D** and **E**) Statistical significance was calculated by one-way ANOVA with Tukey post-hoc test. **, *** and **** denotes p<0.01, 0.001 and 0.0001 in comparison to WT conditions. #### is p<0.0001 in comparison to WT-MIRO1 + MYO19-E→R.

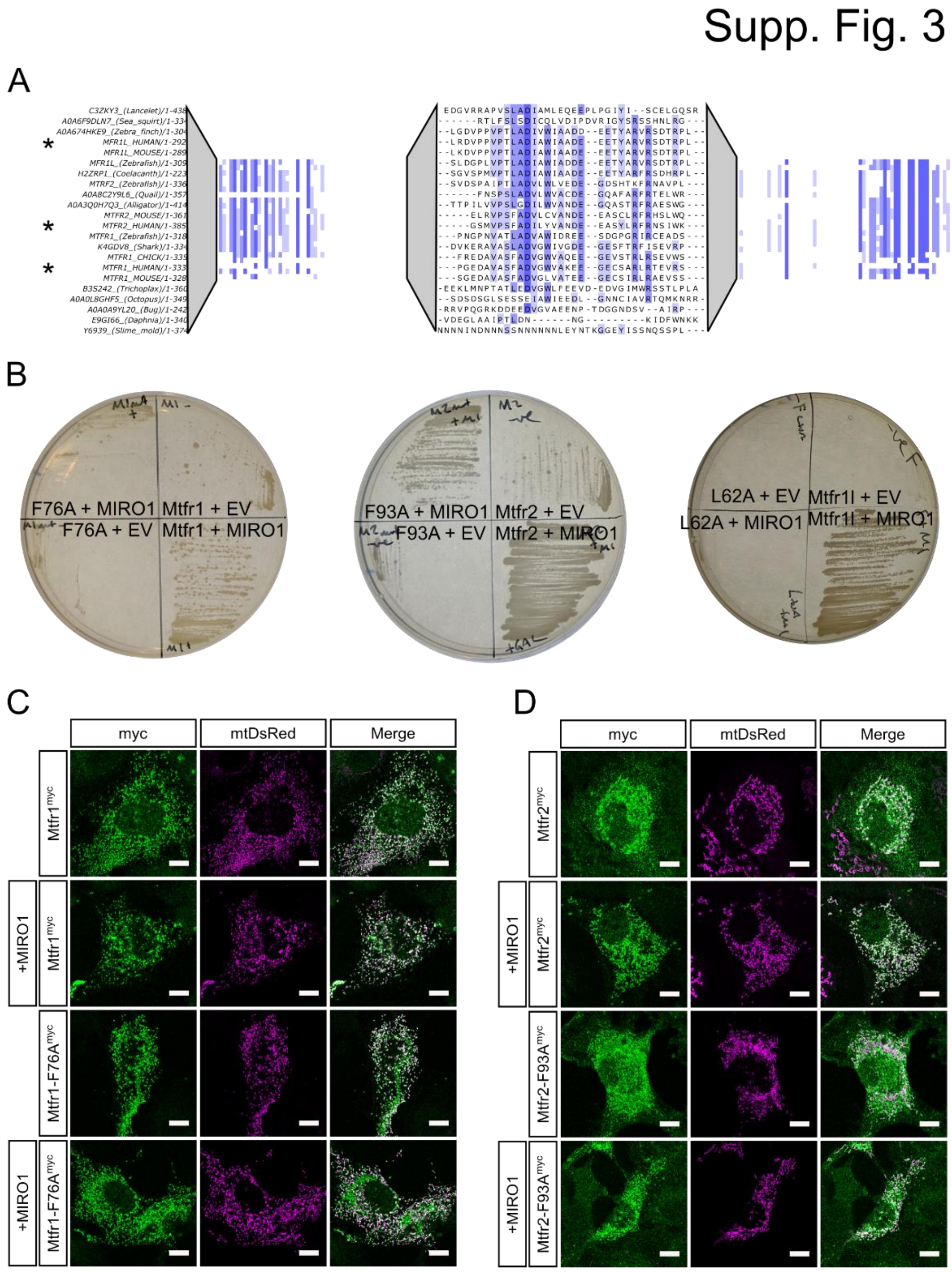


**Fig. S3. Mtfr1/2/1L interact with MIRO1 via conserved motif.** (**A**) Sequence alignment of sequences of MTFR1, MTFR2 and MTFR1L surrounding the Miro-binding motif. (**B**) Yeast two-hybrid growth assay of MIRO1 (prey) with wild-type or point mutants of full-length Mtfr1, Mtfr2 and Mtfr1l. EV means empty vector control. (**C**) Representative images of U2OS Cos7 cells expressing either wild-type or F76A point mutated Mtfr1 (green) both with or without $^{GFP}$MIRO1 overexpression. Mitochondria are stained with mtDsRed (magenta). (D) Representative images of U2OS Cos7 cells expressing either wild-type or F93A point mutated Mtfr2 (green) both with and without $^{GFP}$MIRO1 overexpression. Mitochondria are stained with mtDsRed (magenta). Scale bars depict 10 µm.

### Supp. Fig. 4

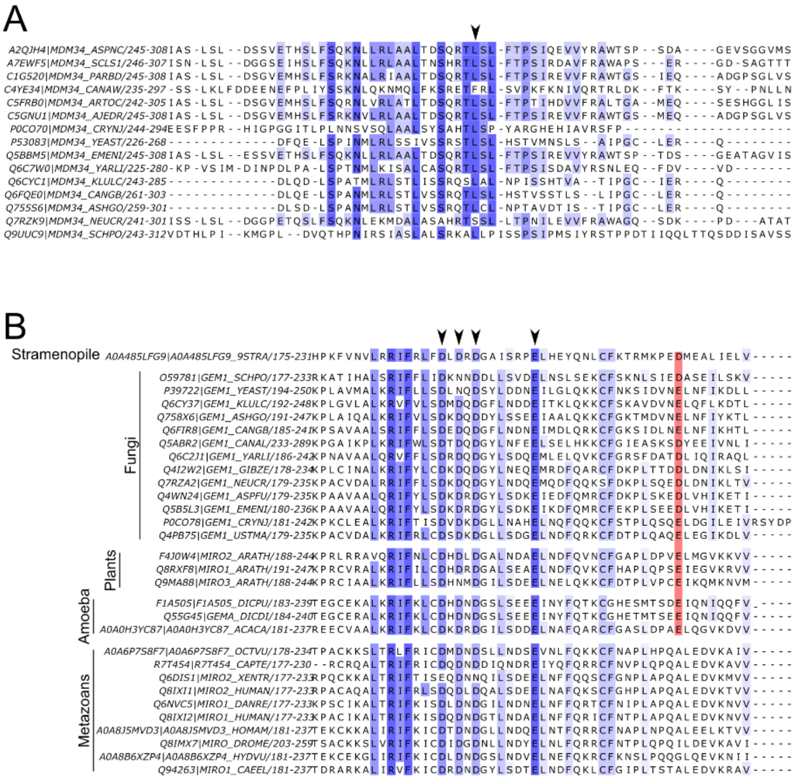


**Fig. S4. Conservation of Gem1 and Mdm34.** (**A**) Sequence alignments of Mdm34. Arrowheads indicate the position of the Leucine inserted in the ELF. (**B**) Sequence alignments of Miro orthologues in fungi, plants, amoeba, and metazoans. Arrowheads indicate the acidic residues coordinating $Ca^{2+}$ in the EF-hand. Red highlights key acidic residue not found in metazoans that is likely required for Mdm34 binding.

Supp. Fig. 5

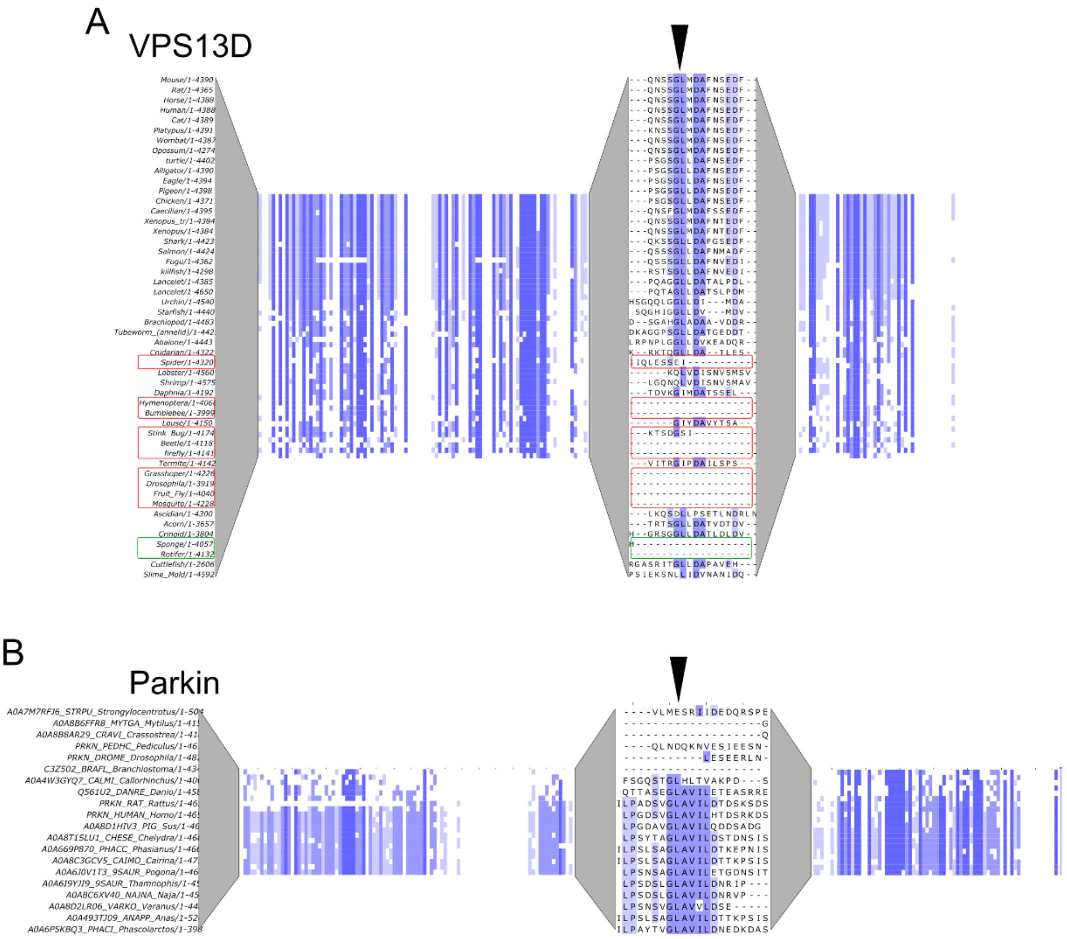


**Fig. S5. Conservation of VPS13D and Parkin Miro binding motifs.** (**A**) Sequence alignments of VPS13D. Red boxes show arthropod sequences that appear to have lost the MBM. The green box shows sequences from primitive animals that are also lacking the MBM (**B**) Sequence alignments of Parkin. Arrowheads point to the conserved leucine residues mutated in this study.

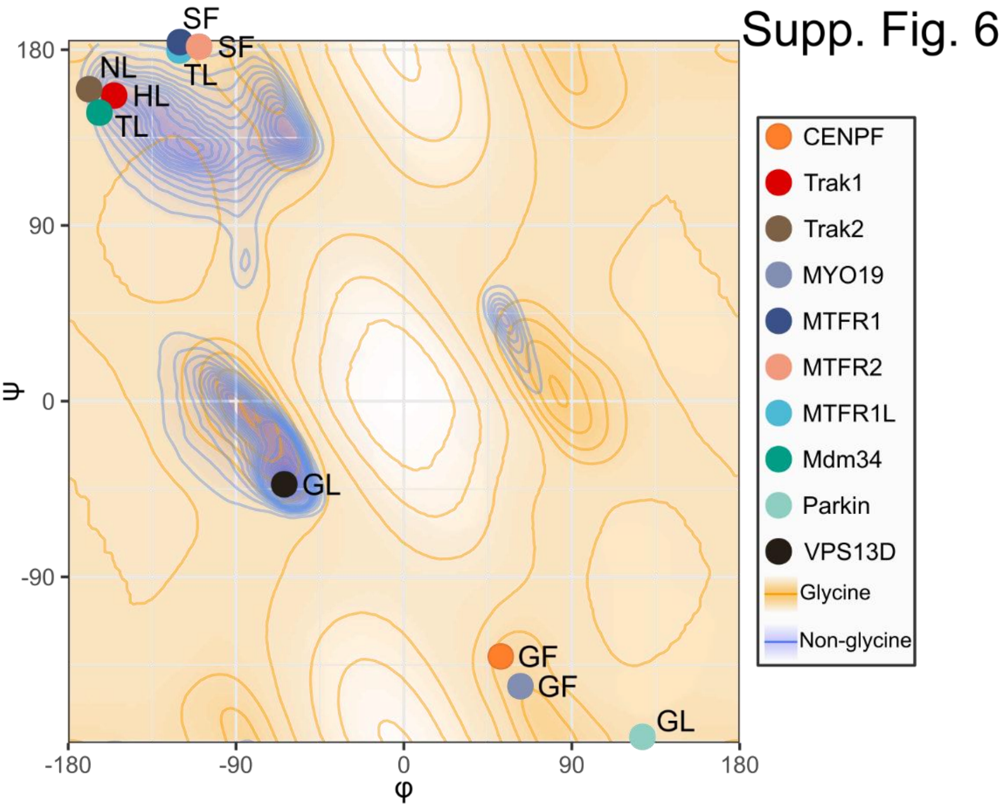


**Fig. S6. Angle requirements of different Miro-binding motifs.** (**A**) Ramachandran plot depicting the bond angles of the residue preceding the ELF-binding F or L from different Miro-binding motifs. Blue and Orange depict the areas tolerating non-glycine and glycine residues respectively.
